# Neutralisation sensitivity of the SARS-CoV-2 BA.2.87.1 variant

**DOI:** 10.1101/2024.03.21.586176

**Authors:** Daniel J. Sheward, Ulrika Marking, Oscar Bladh, Sofia Held, Elise Looyens, Léna Vandenabeele, Sandra Muschiol, Katherina Aguilera, Nina Greilert Norin, Sofia Öling, Michelle Westerberg, Darren Martin, Gunilla B Karlsson Hedestam, Jan Albert, Charlotte Thålin, Ben Murrell

**Affiliations:** Department of Microbiology, Tumor and Cell Biology, Karolinska Institutet, Stockholm, Sweden; Department of Clinical Sciences, Karolinska Institutet Danderyd Hospital, Stockholm, Sweden; Public Health Agency of Sweden; Department of Clinical Microbiology, Karolinska University Hospital, Stockholm, Sweden; Division of Computational Biology, Department of Integrative Biomedical Sciences, Institute of infectious Diseases and Molecular Medicine, University of Cape Town

## Abstract

Against the backdrop of the rapid global takeover and dominance of BA.1/BA.2 and subsequently BA.2.86 lineages, the emergence of a highly divergent SARS-CoV-2 variant warrants characterization and close monitoring. Recently, another such BA.2 descendent, designated BA.2.87.1, was detected in South Africa. Here, we show using spike-pseudotyped viruses that BA.2.87.1 is less resistant to neutralisation by prevailing antibody responses in Sweden than other currently circulating variants such as JN.1. Further we show that a monovalent XBB.1.5-adapted booster enhanced neutralising antibody titers to BA.2.87.1 by almost 4-fold. While BA.2.87.1 may not outcompete other currently-circulating lineages, the repeated emergence and transmission of highly diverged variants suggests that another large antigenic shift, similar to the replacement by Omicron, may be likely in the future.

## MAIN TEXT

The BA.2.86 sublineage of Omicron emerged, phylogenetically rooted near the base of the BA.2 clade indicative of a long period of unobserved evolution prior to its emergence, reminiscent of the initial emergence of Omicron. BA.2.86 had a neutralisation resistance profile that was comparable to other concurrently circulating variants^1^, but its immediate descendent, JN.1, demonstrated rapid growth and swept to global dominance. Recently, yet another highly diverged SARS-CoV-2 variant, designated BA.2.87.1^2^, was reported in South Africa. This novel variant harbours a diverse array of amino acid changes in the receptor binding domain, and two large deletions in the N-terminal domain (Fig A). Against the backdrop of the rapid global takeover and dominance of Omicron (BA.1/BA.2) - and then again BA.2.86 - lineages, this emergence and mutation profile warrants close monitoring and rapid characterization even with only few genomes (N=9) observed to date.

To characterise the relative sensitivity of BA.2.87.1 to current antibody-mediated immunity at the population level, we evaluated BA.2.87.1 neutralisation by sera from blood donations made in Stockholm during week 7 (12th-16th February), 2024. We compared BA.2.87.1 to its parental lineage, BA.2, as well as to XBB.1.5 descendents carrying L455F+F456L (matching GK.3, XBB.1.5.70, and others), and the recently-dominant JN.1 variant. Furthermore, to address the efficacy of the latest vaccines against this variant, we also characterised the responses to BA.2.87.1 in the COMMUNITY cohort^3,4^ prior to and 14 days following a mono-valent XBB.1.5-adapted mRNA vaccine (BioNTech, Mainz, Germany).

BA.2.87.1 was neutralised by sera sampled recently from blood donors with a geometric mean ID_50_ titer (GMT) 2.6x higher than for JN.1 (Fig B). Titers to BA.2.87.1 were only approximately 2.6-times lower than to its parental lineage, BA.2 (GMT of 1121 vs 430) (Supp. Fig. 1). Similarly, in the COMMUNITY cohort, BA.2.87.1 was neutralised by samples taken prior to the XBB.1.5 booster with a GMT approximately 4-times higher than for JN.1 (GMT of 204 vs 53) (Fig B).

**Figure.**
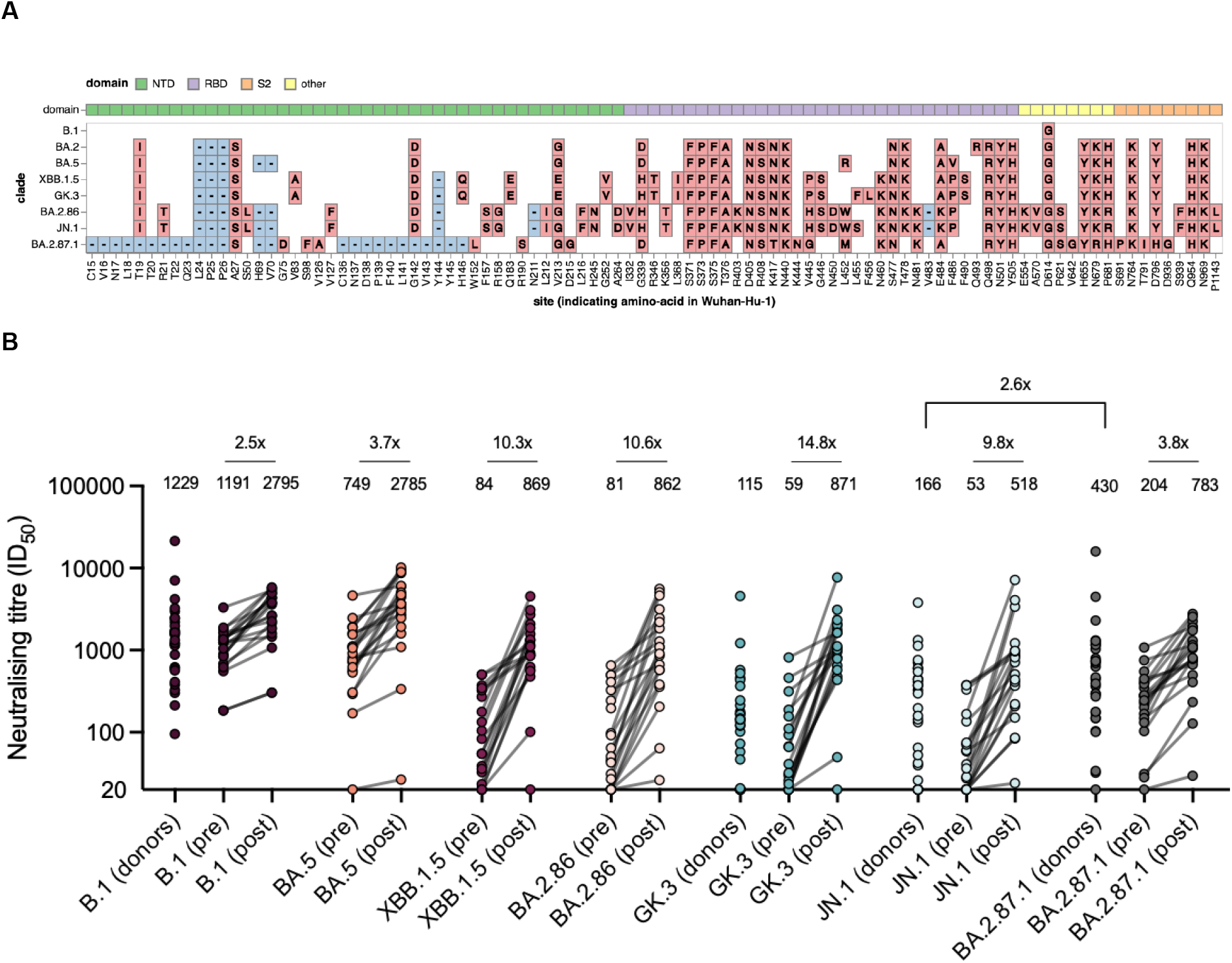
Relative sensitivity of BA.2.87.1 to neutralising antibodies A. A comparison of the amino acid mutations present in the spike protein between multiple variants characterised here, relative to Wu-Hu-1. **B**. Pseudovirus neutralisation by random serum samples (N=30) donated recently (Feb 12th-16th, 2024) in Stockholm, Sweden (donors) and by matched serum samples from the COMMUNITY cohort, taken 0–7 days before (pre) and 14 days after (post) booster vaccination with the monovalent XBB.1.5-adapted BNT162b2 mRNA booster vaccination. The geometric mean ID_50_ titres are summarised above each group, and ID_50_ titres below detection are plotted at the limit of detection (20). ID_50_, half maximal inhibitory dilution. NTD, N-terminal domain. RBD, receptor-binding domain. *** P<0.001.

Neutralising titers to BA.2.87.1 were on average 3.8-times higher 14 days after an XBB.1.5 booster compared to before (Fig C). This is compared to almost 10- to 15-times higher neutralising titers against XBB and BA.2.86 lineages following vaccination, suggesting that the XBB.1.5 vaccine may be relatively less well matched to BA.2.87.1. Nevertheless, neutralising titers to BA.2.87.1 remained higher than those to JN.1, even after the booster (GMT of 783 vs 518).

While BA.2.87.1 was less resistant to neutralisation than other recently circulating variants, it is possible that additional factors affecting the sensitivity to neutralisation are not fully captured in *in vitro* assays. Furthermore, the subset of antibodies mediating the cross-neutralization is not yet resolved. It is possible that additional escape mutations may arise leading to more extensive resistance, as was the case with BA.2.86 leading to the emergence and dominance of JN.1.

While BA.2.87.1 may not outcompete other currently-circulating lineages, the repeated emergence and transmission of highly diverged variants suggests that another large antigenic shift, similar to the replacement by Omicron, may be likely in the future.

## Acknowledgements

pCMV-dR8.2 dvpr was a gift from Bob Weinberg (Addgene plasmid # 8455; http://n2t.net/addgene:8455; RRID:Addgene_8455). pBOBI-FLuc was a gift from David Nemazee (Addgene plasmid # 170674; http://n2t.net/addgene:170674; RRID:Addgene_170674).

We gratefully acknowledge all data contributors, i.e. the Authors and their Originating Laboratories responsible for obtaining the specimens, and their Submitting Laboratories that generated the genetic sequence and metadata and shared via the GISAID Initiative the data on which part of this research is based.

## Supplementary Material

**Supplementary Figure 1.**
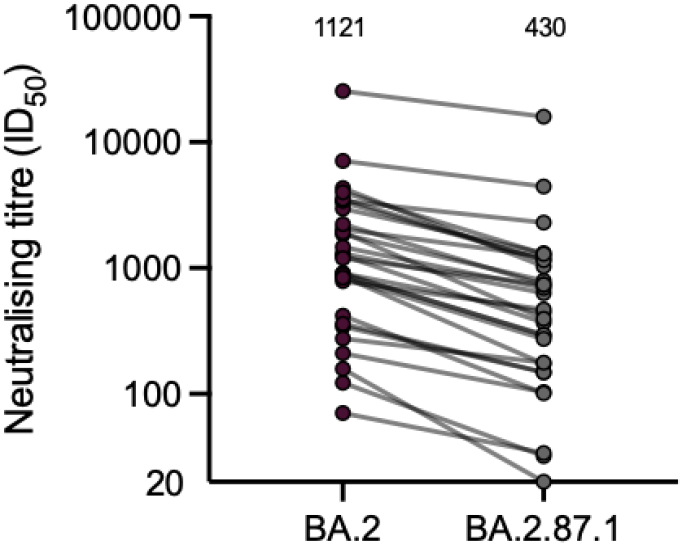
Sensitivity of BA.2.87.1 to neutralisation by random serum (N=30) from blood donated in Stockholm, Sweden between 12th-16th Feb 2024, compared to that for its parental lineage, BA.2. Summarised above each group is the geometric mean ID_50_ titer.

**Supplementary Table 1:**
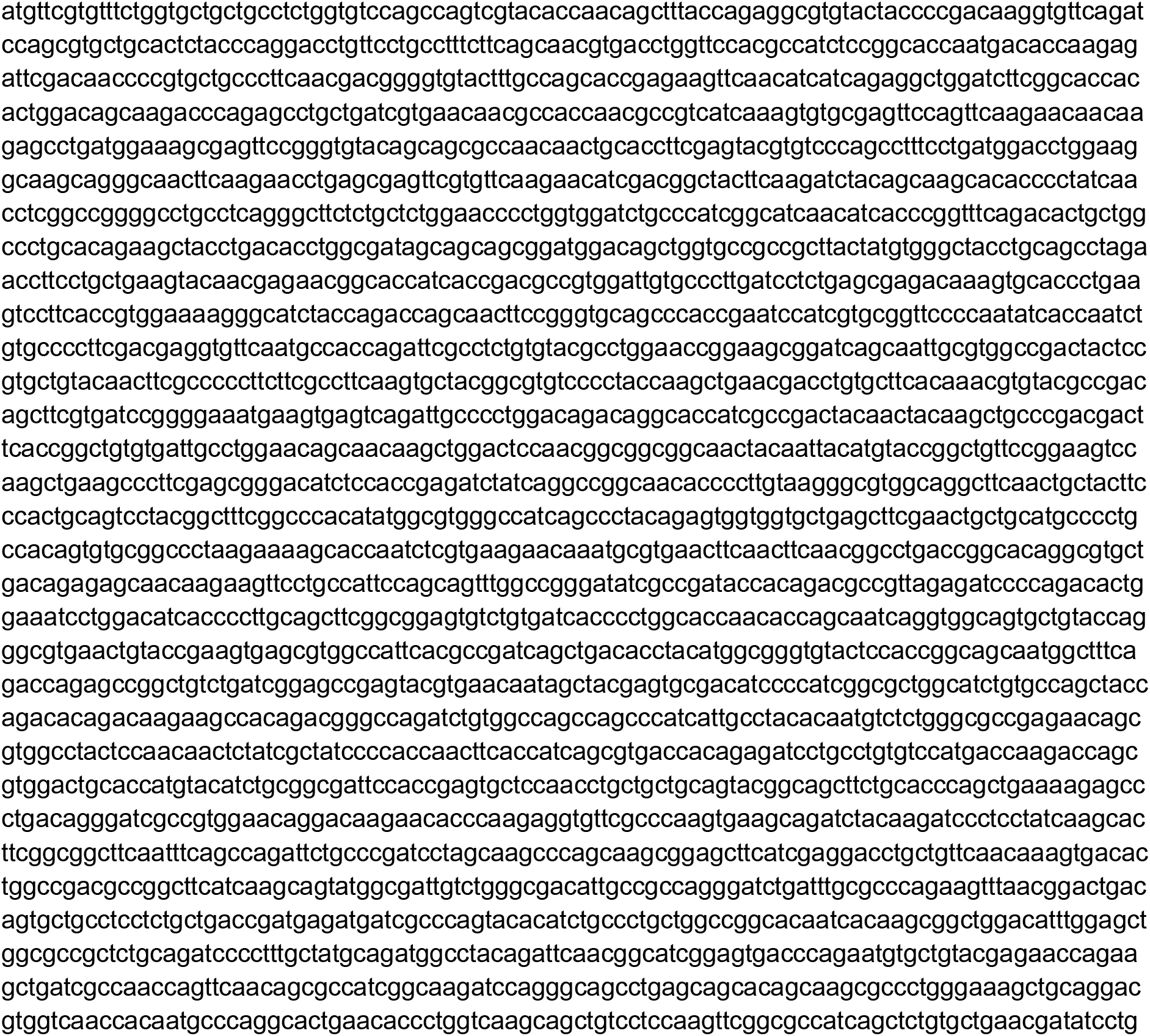
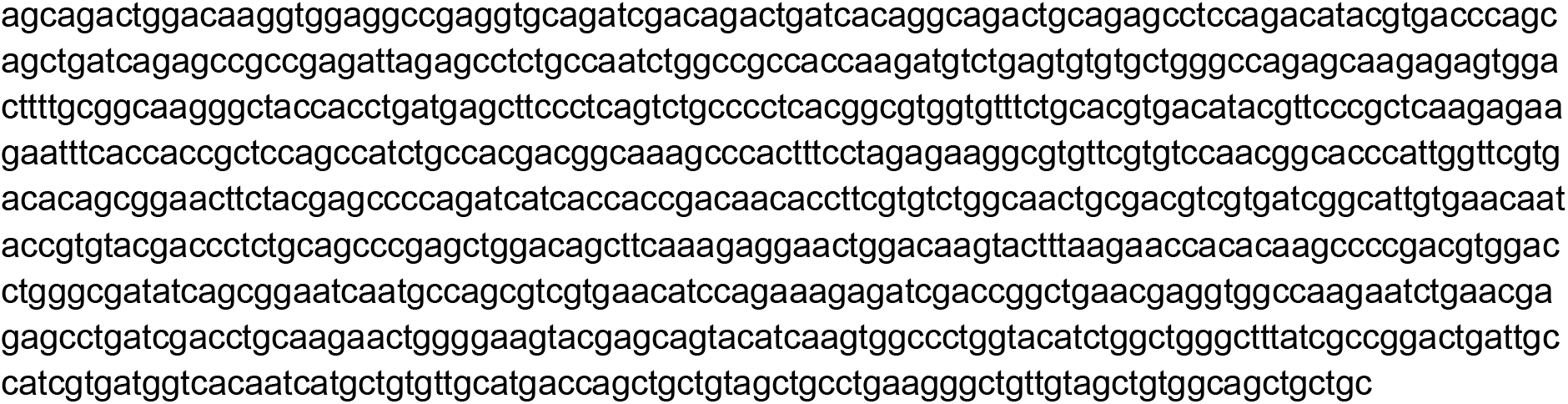
Nucleotide sequence for codon-optimised BA.2.87.1^2^ spike construct:

## Methods

### Cell culture

All cultures were maintained in a humidified 37°C incubator (5% CO2). HEK293T cells (ATCC CRL-3216) and HEK293T-ACE2 cells (stably expressing human ACE2) were cultured in Dulbecco’s Modified Eagle Medium (high glucose, sodium pyruvate) supplemented with 10% fetal bovine serum, 100 units/ml Penicillin, and 100 μg/ml Streptomycin.

### SARS-CoV-2 spike constructs

Codon-optimised BA.2.87.1 spike was synthesised and cloned with Seamless cloning (GenScript). The production of all other codon-optimised spike constructs used here are as previously described^1^. In all constructs, a 19AA cytoplasmic tail truncation is included to promote spike incorporation on pseudoviruses^5^. Plasmids were purified from bacterial cultures using a Qiagen Plasmid Midi kit.

The spike mutation table (Figure A) was generated using the “SARS2-clade-spike-diffs” tool (https://github.com/jbloom/SARS2-clade-spike-diffs).

### Pseudovirus Neutralisation Assay

Pseudovirus neutralisation assays were performed as previously^6^. Briefly, spike-pseudotyped lentiviral particles were generated by the co-transfection of HEK293T cells with respective spike plasmids, an HIV gag-pol packaging plasmid (Addgene #8455), and a transfer plasmid encoding firefly luciferase (Addgene #170674) using polyethylenimine. Neutralisation was assessed in HEK293T-ACE2 cells. Pseudoviruses titrated to produce approximately 100,000 RLU were incubated with 8 serial 3-fold dilutions of serum (beginning at 1:20) for 60 minutes at 37°C in black-walled 96-well plates. 10,000 HEK293T-ACE2 cells were then added to each well, and plates were incubated at 37°C for approximately 48 hours. Luminescence was measured on a GloMax Navigator Luminometer (Promega) using Bright-Glo luciferase substrate (Promega) per the manufacturer’s recommendations. Neutralisation was calculated relative to the average of 8 control wells infected in the absence of antibody. ID_50_ values were calculated in Prism v10 (GraphPad Software) by fitting a four-parameter logistic curve and interpolating the dilution at which there is 50% neutralisation. Assays were repeated at least twice.

### Blood donors

Anonymized serum samples were obtained from random blood donors in Stockholm, as previously described^7^ from week 7 (Feb 12th-16th), 2024. No demographic information is available for these samples. Samples were heat inactivated at 56°C for 45 min prior to use in neutralisation assays.

### COMMUNITY cohort

The COMMUNITY study investigates immune responses to SARS-CoV-2 infection and vaccination among 2,149 healthcare workers enrolled in April 2020 at Danderyd Hospital, Stockholm, Sweden^3^. Vaccination details, including date and type of vaccine, are obtained from the Swedish vaccination registry (VAL Vaccinera), and data on positive SARS-CoV-2 PCR tests are sourced from the Swedish registry for communicable diseases (SmiNet). Clinical and demographic information, including immunocompromising conditions or treatments and information about positive rapid diagnostic tests (RDT), are collected via a smartphone application-based questionnaire during each follow-up visit.

In this sub-study, we investigated serological responses before and after a monovalent XBB.1.5-adapted BNT162b2 mRNA booster dose (30 μg) in 24 immunocompetent participants without intervening infections, as previously described^4^. Samples were heat inactivated at 56°C for 45 min prior to use in neutralisation assays.

## Ethical Statement

The study was approved by the Swedish Ethical Review Authority (dnr 2020-01653) and conducted in accordance with the declaration of Helsinki. Written informed consent was obtained from all COMMUNITY study participants. The blood donor samples were anonymized and therefore could be analysed without informed consent from the donors, as per advisory statement 2020–01807 from the Swedish Ethical Review Authority and the Swedish Ethics Review Act.

## Statistical analysis

Neutralising titres between groups were compared using a Wilcoxon matched-pairs signed rank test in Prism v10 (GraphPad Software).

## Reagent availability

The BA.2.87.1 spike expression plasmid used here is available upon request.

## Author contributions

Conceptualization, D.J.S., D.M., C.T., B.M.; Formal analysis, D.J.S, B.M.; Conducted the assays, D.J.S., S.H., E.A.L.L., L.V. S.Ö., M.W.; Designed the methodology, D.J.S; Responsible for figures and tables, D.J.S, B.M.; Resources, U.M., O.B., S.M., D.M., G.B.K.H., J.A., C.T., B.M.; Oversaw the study, D.J.S., C.T., B.M; Funding Acquisition, G.B.K.H., J.A., C.T., B.M., Writing – original draft, D.J.S., B.M.; Writing – review & editing, D.J.S., G.B.K.H., J.A., C.T., B.M.

D.J.S, C.T., and B.M. were responsible for the decision to submit the manuscript for publication.

